# QTL mapping in field plant populations reveals a genetic basis for frequency- and spatially-specific fungal pathosystem resistance

**DOI:** 10.1101/2024.06.03.597112

**Authors:** Patrycja Baraniecka, Klaus Gase, Maitree Pradhan, Ian T. Baldwin, Erica McGale, Henrique F. Valim

## Abstract

Fungal pathogens pose significant challenges to agro-ecosystem productivity. The wild tobacco, *Nicotiana attenuata*, has been grown for over two decades at an experimental field station in its native habitat, leading to the emergence of a high-mortality sudden wilt disease caused by a *Fusarium-Alternaria* pathosystem. By using an Advanced Intercross Recombinant Inbred Line (AI-RIL) mapping population of *N. attenuata* planted in the infected field site, we found two significant loci associated with plant susceptibility to the fungi. A functional characterization of several genes in these loci identified *RLXL* (intracellular ribonuclease LX-like) as an important factor underlying plant pathogen resistance. Virus-induced silencing of *RLXL* reduced leaf wilting in plants inoculated with an *in vitro* culture of *Fusarium* species. Assessing the significance of the *RLXL*-associated allele in mixed field populations indicated that, among 4-plant subpopulations, those harboring a single plant with the *RLXL*-deficiency allele exhibited the highest survival rates. Within these populations, a living *RLXL*-deficient plant improved the survival of *RLXL-*producing plants located diagonally, while the mortality of the adjacent plants remained as high as in all other subpopulations. Taken together, these findings provide evidence for the genetic basis for a frequency- and spatially-dependent population pathogen resistance mechanism.

**Significance statement:** Plant pathogen resistance studies predominantly focus on single genes that reduce pathogenicity in individual plants, aiming to apply these findings to agricultural monocultures. On the other hand, ecologists have observed for decades that greater diversity drives plant population resistance and resilience to pathogens. More studies are needed to identify and characterize loci with positive effects conferred through their frequency in plant populations. We combine quantitative genetics, molecular techniques, and ecologically-informed mixed field populations to identify a novel intracellular ribonuclease LX-like (*RLXL*) gene with a frequency- and position-dependent effect for plant resistance. To our knowledge, this is the first detailed link between plant population protection and various percentages of plants with an allele representing *RLXL* presence or absence.

## Introduction

*Nicotiana attenuata* is an annual, wild coyote tobacco species that germinates after wild fires from long-lived seedbanks. This characteristic contributes to the development of genetically diverse plant communities that contrast to agricultural monocultures ^1^. A rapidly spreading sudden wilt disease, identified as a *Fusarium-Alternaria* pathosystem ^2^, was first observed within an experimental field plot in the Great Basin Desert. Repeated, near-monoculture plantings of *N. attenuata* eliminated the extensive genetic diversity usually found in natural populations, leading to pathogen accumulation in the soil and increased plant mortality rates ^3^. Infected plants wilted, turned black, and eventually collapsed, resulting in the loss of over half of the plants in the affected plot ^3^. The significant impact of genetic diversity on plant survival was highlighted by a comparable disease outbreak in the neighboring natural population of *N. attenuata*. Unlike the experimental plantation, where the disease persisted, many plants in the natural population recovered within the same season with no lasting impact of the disease ^2^.

Disease incidence and plant mortality in the experimental field were shown to be reduced by inoculation with a mixture of native bacterial isolates ^3^. Subsequent analysis of the molecular components involved in related plant resistance showed that jasmonic acid (JA) and O-acyl sugars (O-AS) play an important role in mediating plant response to the pathosystem ^4, 5^. Recently, pathogen-induced JA signaling responses were also shown to be modulated by AGO4 ^6^. However, to date, the genetic and molecular mechanisms enabling the natural populations of *N. attenuata* to quickly counteract the dynamic pathosystem and prevent long-lasting effects remain unidentified.

Here, we describe the genetic factors underlying the resistance of *N. attenuata* to the devastating effects of *Fusarium-Alternaria* pathosystem and examine their contribution to the survival of individuals within population in the context of genetic diversity. By utilizing an Advanced Intercross Recombinant Inbred Line (AI-RIL) mapping population and a novel resistance score calculated for each AI-RIL planted in the field, we identified two significant loci associated with individual plant mortality. Further analyses led to the identification of intracellular ribonuclease LX-like (*RLXL*) as a main factor linked to differences in plant survival. Virus-induced silencing of *RLXL* significantly reduced leaf wilting, which is the main symptom of the pathogen infection. We investigated the ecological function of this gene, hypothesizing that *RLXL* may contribute to population resistance in a manner dependent on both gene frequency and spatial distribution.

## Results

### QTL mapping on the resistance of N. attenuata to a Fusarium-Alternaria pathosystem in the field yields two significant loci

To identify the genetic basis underlying the resistance of *N. attenuata* to its native *Fusarium-Alternaria* pathosystem, we performed quantitative trait locus (QTL) mapping using survival data on an advanced intercross - recombinant inbred line (AI-RIL) population planted in a field site with a persistent and abundant pathosystem ^3, 7^. As the presence-absence trait of survival cannot be mapped on directly, each AI-RIL line was assigned a resistance score (RS; Fig. 1a-d). This approach not only provided quantitative data, but also allowed us to normalize RS to a control plant in each four-plant subpopulation and account for differences in pathosystem load across the field site (Fig. 1c). Within the 90% confidence interval we found two loci linked to the plant RS (Fig. 1e). Within the co-inherited regions to these loci, we identified several candidate genes (Table S1). Two particularly promising candidates, intracellular ribonuclease LX-like (*RLXL)* and ABC transporter G family member 23-like (*ABCG23*), were selected based on their functional annotation, expression in tissues relevant to pathogen modes of action, and an extensive literature review.

**Figure 1:**
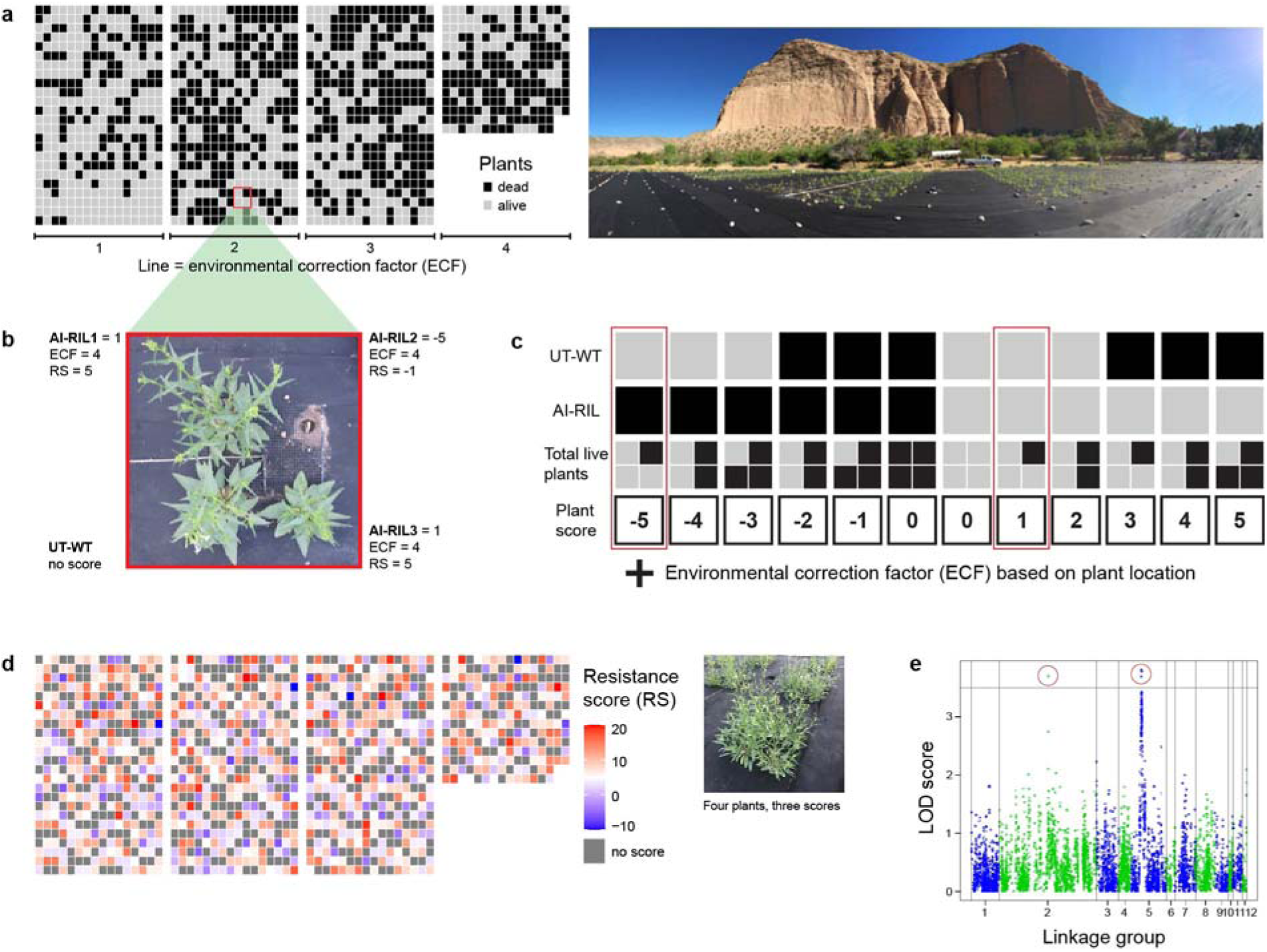
QTL mapping on the resistance of *N. attenuata* plants to a *Fusarium-Alternaria* pathosystem in the field yields two significant loci. (a) A mortality map (left) of the advanced intercross-recombinant inbred line (AI-RIL) population interspersed with UT-WT control plants grown at the field site at Lytle Ranch Preserve, UT, USA, in 2017 with a picture of the field plot (right). (b) All plants were arranged in four-plant subpopulations around a single drip irrigation emitter (pictured) with three randomly chosen AI-RIL lines and a UT-WT control. (c) The system used to determine the resistance score of each AI-RIL line. The scores for the AI-RIL lines shown in (b) are highlighted in red. These would then be adjusted by the environmental correction factor (ECF) to produce the final resistance score (RS), as shown in (b). Grey and black squares indicate live and dead plants, respectively in both (a) and (b). (d) The sum of RS for the four replicates of each AI-RIL is displayed at each location of that AI-RIL line. AI-RIL lines with less than four replicates were removed from the analysis and are shown in gray. An example four-plant population with three scores (as UT-WT does not receive a score) is shown on the left. (e) Quantitative trail locus (QTL) mapping of the RS from (d) resulted in two significant groups of loci (above the 95% confidence interval of 3.5 LOD; red circles). LOD: log of odds.

### Changes in the transcript abundance dependent on RLXL-associated allele correlate with plant survival

To validate the field results and further investigate the target genes, we conducted a second field experiment in the following year. The two founder lines of the AI-RIL population, UT- and AZ-WT, as well as AI-RIL 106A, which had one of the highest RS in the original experiment, were planted in the same field plot. At the *RLXL*-associated locus, line 106A has the same allele as AZ-WT and the same allele as UT-WT at the *ABCG23*-associated locus. Only 27% of UT-WT plants survived to the end of the experiment (Fig. 2a). In contrast, AZ-WT and line 106A showed significantly higher survival rates, with 72% (*P* = 0.0151) and 52% (*P* = 0.0303) of plants surviving until the final harvest, respectively. These results were further verified in an *in vitro* seedling experiment where the same lines were inoculated with lab cultures of either *Fusarium brachygibbosum* or *Alternaria* sp. (Fig. 2b,c). AZ-WT and 106A seedlings showed significantly higher survival in the face of both pathogens relative to UT-WT (*P* < 0.0001 for both lines, for both pathogens), indicating that the *RLXL*-associated allele shared between these two lines is more likely to explain the observed resistance effect.

**Figure 2:**
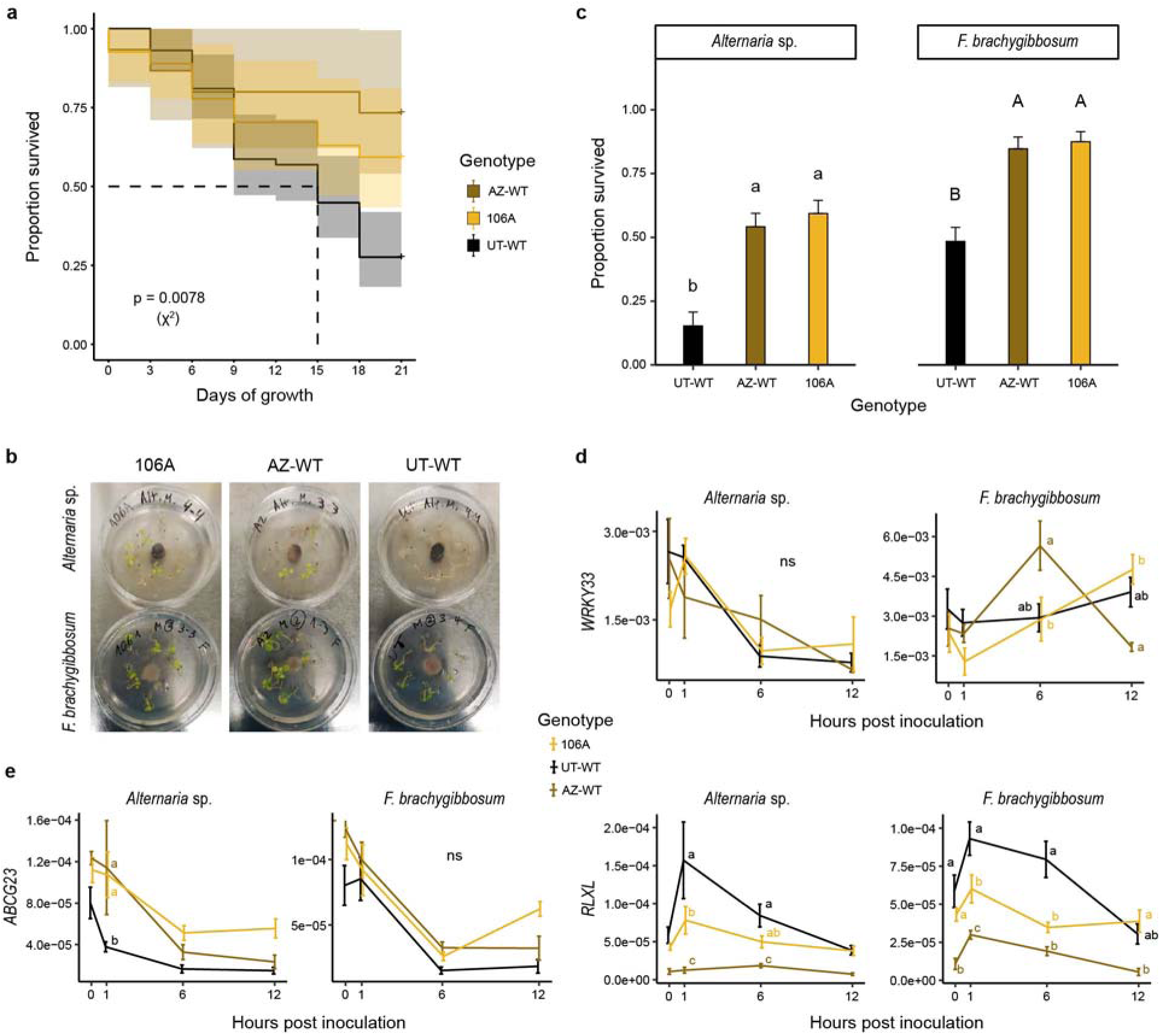
Survival of AZ-WT and line 106A in the field and *in vitro* correlates with changes in the abundance of *RLXL* transcripts. (a) Proportion survived (± CI, n = 15-58) of the focal genotypes AZ-WT, 106A and UT-WT planted at the field site at Lytle Ranch Preserve, UT, USA during the 2018 field season based on a multivariate Cox regression analysis. (b) Experimental design for an *in vitro* seedling test: plates inoculated with either *Alternaria* sp. or *F. brachygibbosum* fungal plugs placed in equal distance from each of the eight seedlings. (c) Proportion survived (± SE, n = 6 – 9) of seedlings after inoculation with either *Alternaria* sp. or *F. brachygibbosum*. Relative transcript abundance of (d) a pathogen response related transcription factor (positive control) and (e) a subset of genes within linkage disequilibrium of the significant QTL (full names and NIAT identification numbers are shown in Table S1). Small letters indicate statistically significant differences between the genotypes within one timepoint based on ANOVA followed by Tukey adjusted pairwise contrasts.

We also measured transcript abundance of *RLXL* and *ABCG23* in seedlings at 0-, 1-, 6- and 12-hours post inoculation (hpi), employing the transcription factor *WRKY33* as a positive control given its previously characterized role in plant pathogen resistance (Fig. 2d,e). The conserved accumulation of *WRKY33* transcript abundance among the three lines was confirmed after inoculation with *Alternaria* sp., while it was only true up to 1 hpi with *F. brachygibbosum* (Fig. 2d). Therefore, to evaluate the effect of the *Fusarium* species on the transcript abundance of *ABCG23* and *RLXL,* we focused only on the early changes (Fig. 2d). *F. brachygibbosum* inoculation caused no differences in the accumulation of *ABCG23* transcripts between the three lines, while inoculation with *Alternaria* sp. resulted in rapid and significant increases in AZ-WT and 106A seedlings relative to UT-WT. In contrast, the transcript abundance of *RLXL* was significantly lower in AZ-WT and 106A compared to UT-WT seedlings after inoculation with both pathogens (Fig. 2e).

Due to the previously described role of jasmonate (JA) and salicylic acid (SA) in mediating the response of *N. attenuata* to its native pathogens ^4^, we investigated whether the changes in transcript abundance of genes related to these plant hormone pathways responded to inoculation with *F. brachygibbosum* and *Alternaria* sp. in a manner similar to *ABCG23* or *RLXL* (Fig. S1). Among the tested genes, only *PR1* (pathogenesis-related 1), which is known to be involved in SA-mediated pathogen defense ^8^, showed differences in transcript accumulation between the different lines, although only before pathogen inoculation. None of the tested genes showed differences among the three lines by 1 hpi, which contradicted the potential downstream connection to the candidate genes.

### RLXL transcript accumulation reflects survival among N. attenuata natural accessions

We investigated the extent of the variation in *RLXL* expression and its impact on seedling survival after the inoculation with *F. brachygibbosum* in an independent set of *N. attenuata* natural accessions, compared to UT- and AZ-WT controls (Fig. 3). Prior to inoculation, the highest *RLXL* transcript abundance was observed in line P108 and UT-WT, while it was significantly lower in AZ-WT and line P370 (Fig. 3a). The lowest *RLXL* transcript abundance in these accessions corresponded to the highest seedling survival rates whereas higher accumulation of *RLXL* transcript decreased the survival by as much as 50% (Fig. 3b). Notably, we did not observe a negative correlation between *RLXL* transcript abundance and seedling survival across different accessions following *Alternaria* sp. (Fig. S2).

**Figure 3:**
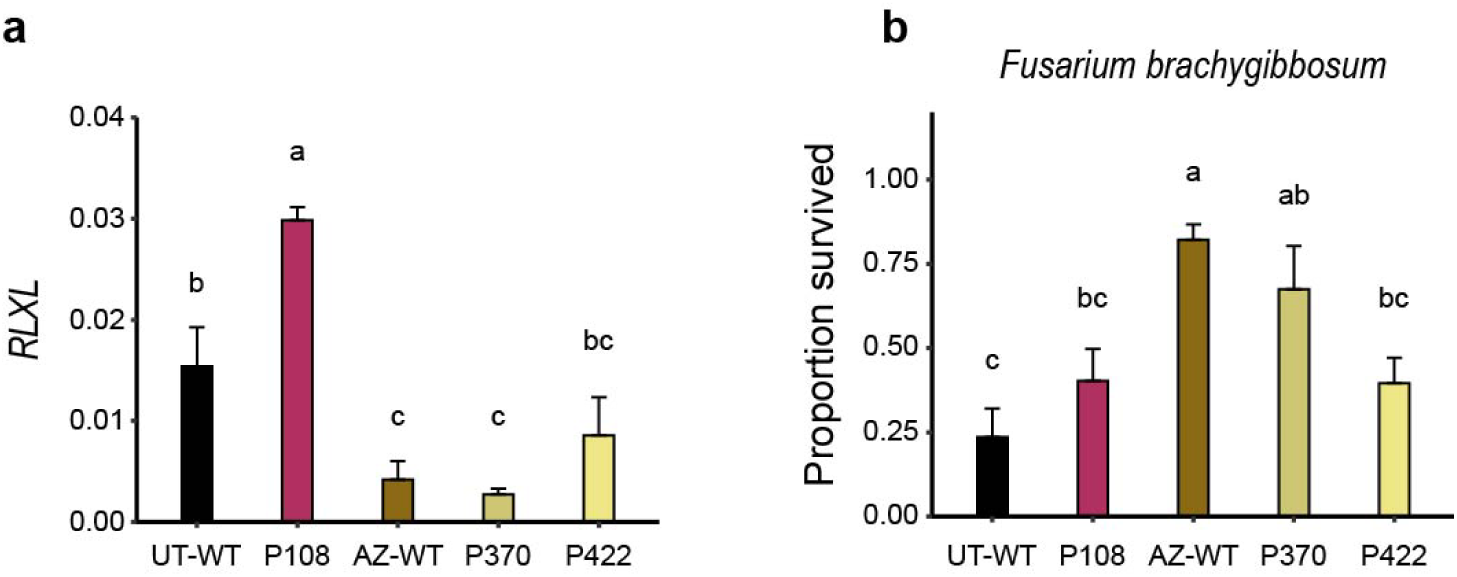
Seedling survival of natural *N. attenuata* accessions is linked to the *RLXL* transcript abundance. (a) Relative transcript abundance of *RLXL* (mean ± SE, n = 3-4) in different natural accessions of *N. attenuata*. (b) Proportion survived (± SE, n = 6-9 per accession) of seedlings inoculated with *Fusarium brachygibbosum*. Small letters indicate statistically significant differences based on ANOVA followed by Tukey adjusted pairwise contrasts, p < 0.05.

### Silencing of RLXL reduces leaf wilting symptoms caused by F. brachygibbosum

To directly examine the function of *RLXL* in *N. attenuata* resistance to *F. brachygibbosum*, we silenced the gene in UT-WT and P108 plants (both displaying high initial accumulation of *RLXL* transcript, Fig. 3) using Tobacco Rattle Virus-Induced Gene Silencing (VIGS; Fig. 4a,b). At 21 days after VIGS (dav) the *pTV::PDS5* VIGS positive control plants showed clear signs of bleaching in newly grown leaves (Fig. 4a), indicating successful silencing and relevant aboveground tissues to sample. Silencing efficiency of *RLXL* in roots was confirmed by qRT-PCR, which showed that the gene expression was significantly lower in both UT-WT::*RLXL* and P108::*RLXL* silenced plants (∼60% reduction) compared to their empty vector (*EV*) controls (UT::*EV* and P108::*EV*, Fig. 4c).

**Figure 4:**
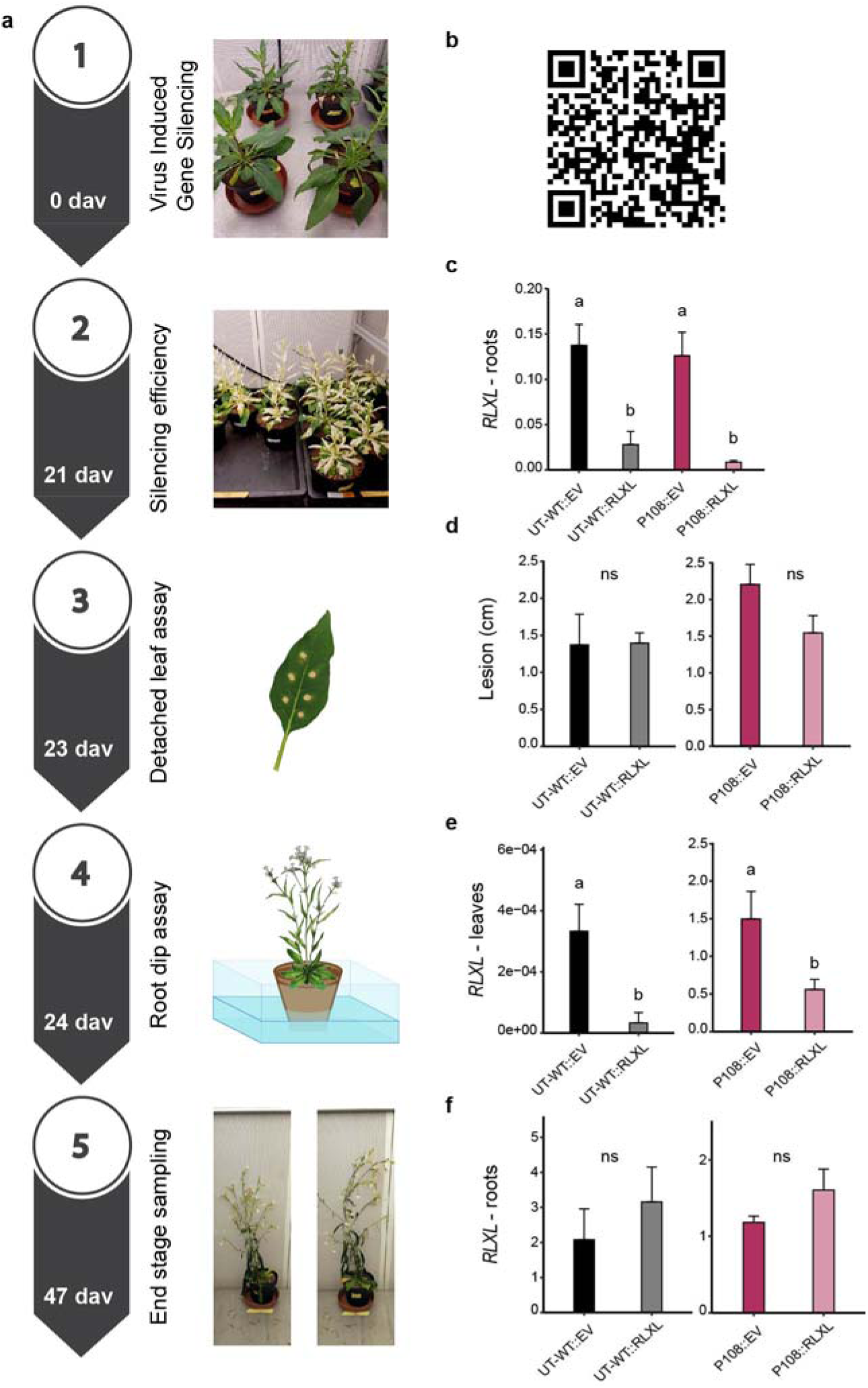
Virus-Induced Gene Silencing (VIGS) of *RLXL* reduces transcript abundance in leaves and roots for pathogenesis studies. (a) Timing of the VIGS experiment and assays performed at each day after VIGS (dav). (b) QR code linked to a video demonstrating the VIGS inoculation. (c) Relative *RLXL* transcript abundance in the roots of a subset of plants destructively sampled 21 dav. (d) Lesion size (6 lesions per leaf) on detached leaves six days post inoculation (dpi) with *F. brachygibbosum* leaf inoculation assay. Relative *RLXL* transcript abundance in (e) leaves and (f) the root-dip inoculated roots at the final harvest of the VIGS plants (59 days after germination, 47 dav, and 23 dpi). Lowercase letters indicate statistically significant differences based on ANOVA followed by Tukey adjusted pairwise contrasts within each accession (± SE, n = 3-4).

A detached leaf assay was performed at 23 dav on all VIGS plants to evaluate fungal pathogenicity. No differences in the size of the lesions were observed between the silenced lines and controls (Fig. 4d). It has been shown previously that the pathogen enters the plant through the roots, up to the root-shoot junction, causing severe stem and leaf wilting ^2–4^. Therefore, to investigate the effect of *RLXL* silencing on plant pathogen responses *in vivo* and in an ecologically relevant manner, fully-grown VIGS plants were inoculated with *F. brachygibbosum via* a root dip assay at 24 dav. The analysis of *RLXL* transcript abundance in both leaves and roots of these plants at the end of the VIGS experiment (23 days post inoculation, dpi and 47 dav) revealed sustained silencing in leaves, while in roots the difference between -::*RLXL* and -::*EV* was no longer significant (Fig. 4e,f), consistent with the decrease in *RLXL* transcript abundance in all lines after pathogen inoculation in seedlings (Fig. 2e) and further indicating the tissue specific nature of *RLXL*-mediated responses. We also monitored leaf wilting in the VIGS plants after root dip inoculation (Fig. 5). Up to 15dpi, there were no significant differences in the percentage of wilting leaves, and relative growth rate (RGR) observed between the -::*RLXL* and their -::*EV* controls, although *-::RLXL* plants showed a strong tendency for reduced wilting relative to -::*EV* controls (Fig. 5b,c). At 23 dpi, leaf wilting was at least marginally significantly reduced in all *RLXL-*silenced plants, as measured by leaf angle of the remaining leaves (Fig. 5d), with less distinct differences in the P108::*RLXL*, possibly due to the differences in leaf size (Fig. 5e).

**Figure 5:**
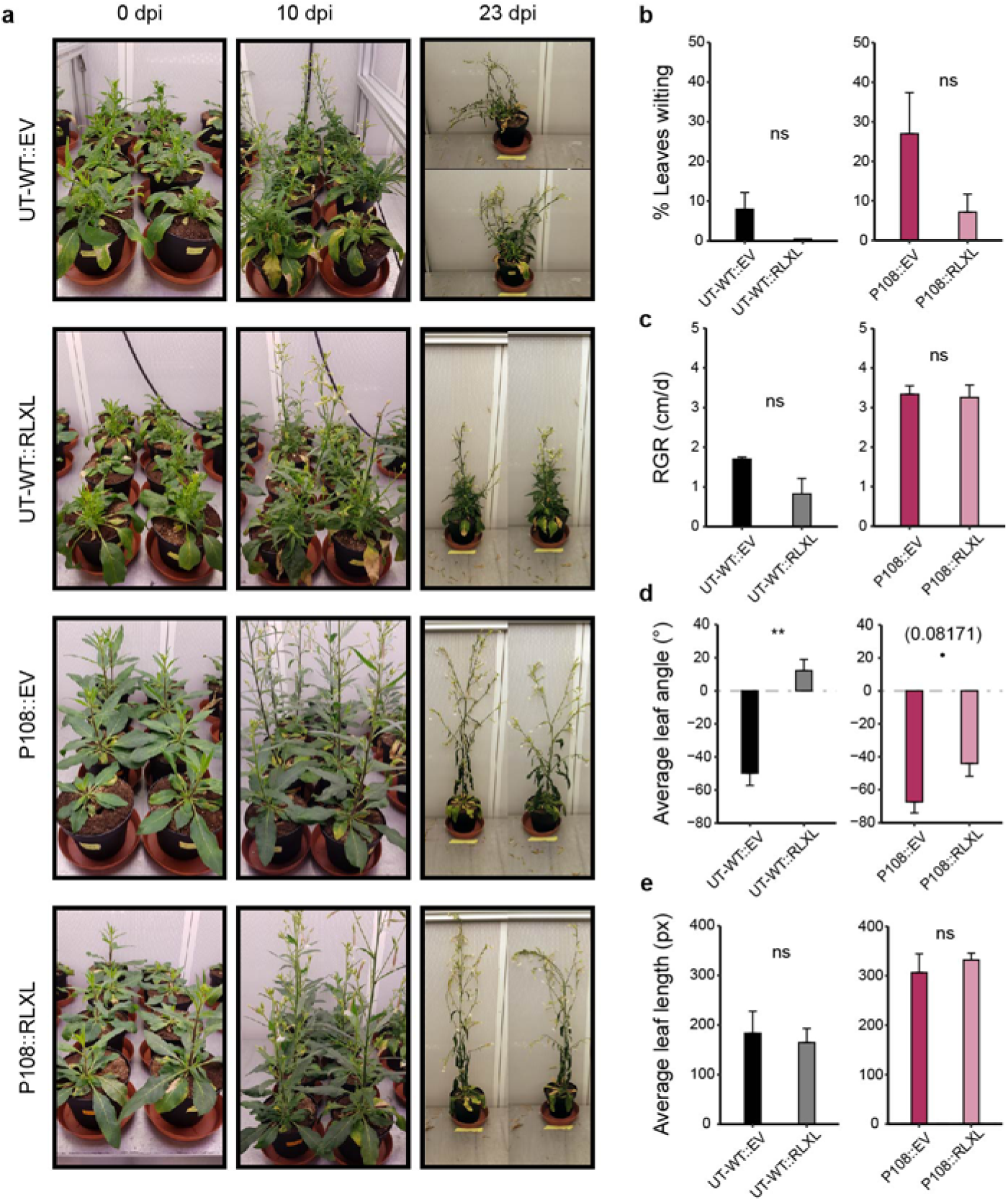
Silencing of *RLXL* reduces leaf wilting caused by the *F. brachygibbosum* root dip inoculation in *N. attenuata*. (a) VIGS plants pictured at 0-, 10- and 23 days post inoculation (dpi). (b) Percentage of wilting leaves and (c) the relative stalk growth rate (RGR) of VIGS plants at 15 dpi. (d) Average leaf angle and (e) leaf length at 23 dpi. Lowercase letters indicate statistically significant differences based on ANOVA followed by Tukey adjusted pairwise contrasts within each accession (± SE, n = 3-4).

### Effect of RLXL-deficient allele on plant survival depends on frequency and position within field populations

To assess the impact of *RLXL* on population survival from an ecological perspective, we investigated the varying frequencies of *RLXL*-deficient allele within the randomized AI-RIL population design of the original field experiment (Figs 1b, 6). First, we examined the number of surviving plants in populations with at least one dead plant and varying numbers of *RLXL­*-deficient (found in AZ-WT accession – the ‘A’ allele) alleles across field locations that displayed increasing disease intensity (Figs. 1a, 6b). We observed the highest plant survival in populations with one A allele across the field locations with low and medium disease intensity (Fig. 6b).

**Figure 6:**
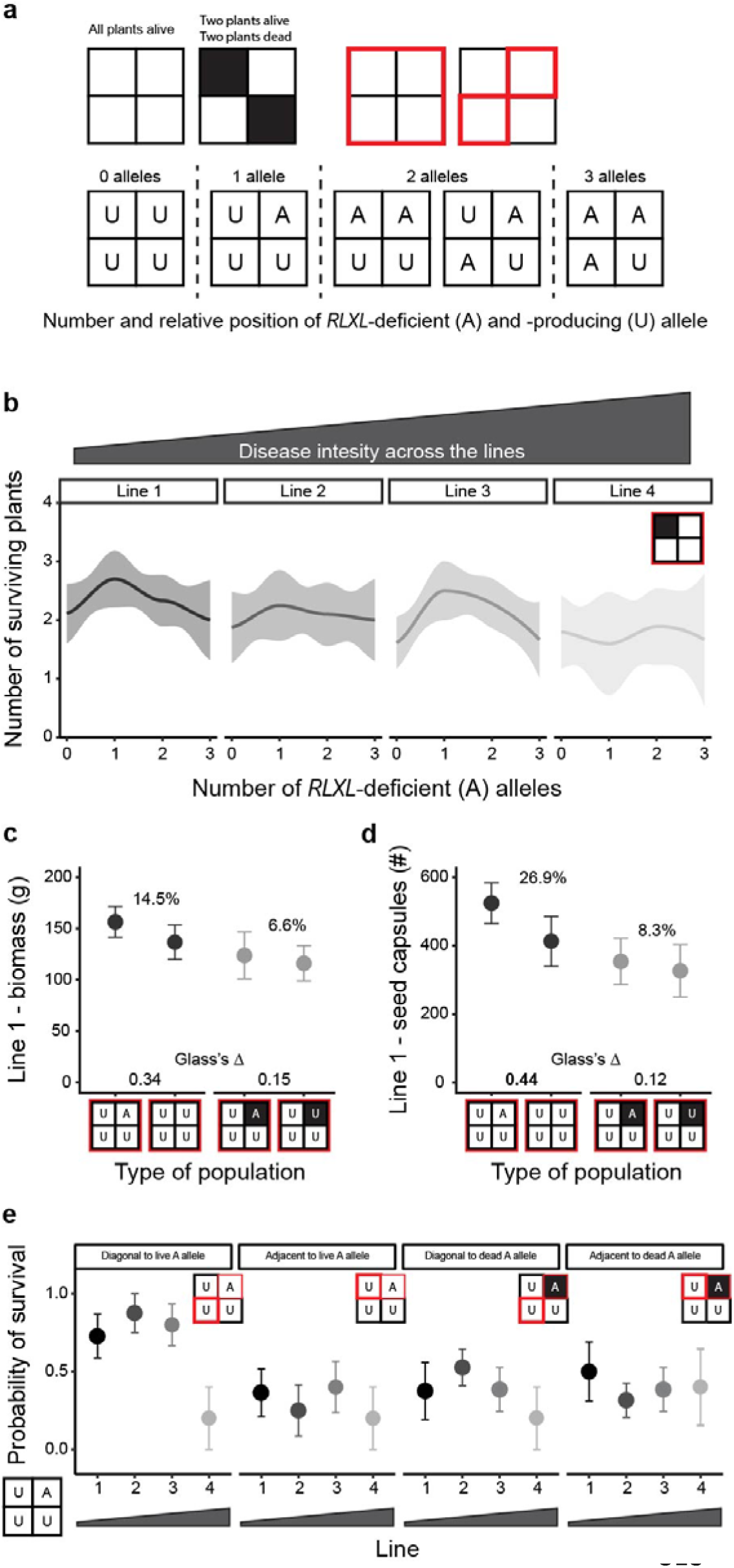
Effect of the allele associated with *RLXL*-deficiency (A) on plant survival depends on its frequency and position. (a) Schematic representation of different four-plant populations (Lytle Ranch Preserve, UT, USA, 2017) with live (white) *vs.* dead (black) plants. All four squares are highlighted in red when the whole population is considered for an analysis. Individual squares are highlighted in red when the data originates from those particular plants with relevance to their position to another plant (top). Possible combinations of number and relative position of *RLXL*-deficient (found in AZ-WT accession – the ‘A’ allele) and -producing (found in UT-WT accession – the ‘U’ allele) plants are shown (bottom). (b) Number of plants surviving (± SE) in individual four-plant populations with different frequencies of A allele plants when at least one plant is dead (regardless of its *RLXL*-associated allele), across the four field locations with increasing disease intensity (top grey triangle). (c) Biomass accumulation (± SE, n = 8-12) and (d) number of ripe and unripe seed capsules (± SE, n = 8-12) in populations with one or no A compared when all four plants were alive (left, black) or at least one plant died (right, grey). Glass’s Δ and percentage difference is reported for each pair, medium effect size is indicated in bold. Detailed statistical information for (c) and (d) is shown in Table S2. (e) Probability of survival of individual plants growing either diagonally or adjacent to a plant with the A allele only in populations with a total of one A allele plant (ANOVA_position_ = 0.007).

Subsequently, we investigated the productivity of *RLXL*-diverse populations in the face of disease by measuring plant biomass and reproductive correlates (Fig. 6c,d). In order to ensure sufficiently high numbers of replicates of each type of subpopulation, we compared populations with no A alleles and one A allele only from the field location with the lowest disease intensity. In all subpopulations with the A-plant alive and otherwise all levels of plant survival allowed, we observed 14.5% more biomass than in comparable populations with only *RLXL*-producing alleles (found in UT-WT accession – the ‘U’ allele). In contrast, only 6.6% more biomass was observed when the one A-plant was dead when compared to U-only populations with one dead plant (Fig. 6c). For reproductive correlates, the effect of the live A-plant was even more dramatic, with 26.9% more production when the A-plant was alive *versus* 8.3% when it was dead, as compared to U-only subpopulations (Fig. 6d).

Additionally, we questioned whether spatial distribution may play a role on the positive effect of the single A-plant on population survival and productivity. We analyzed the probability of survival for individual plants growing either diagonally or adjacent to a single A-plant, when that plant was either dead or alive across the field locations with increasing disease intensity (Fig. 6e). Plants grown diagonally to a plant with an A allele had a significant increase in the probability of survival when compared to plants adjacent to the A allele, but only when the A allele plant was alive.

## Discussion

QTL mapping on the resistance score calculated for each line from the AI-RIL mapping population and the subsequent analysis of the genomic regions corresponding to two significant QTLs led to the identification of two promising candidate genes: *RLXL*, and the ABC transporter G family member 23-like (*ABCG23*). ABC transporters have long been implicated in plant pathogen defense, and there is considerable evidence that *ABCG* genes in plants and pathogens have co-evolved to secrete secondary metabolites involved in the plant-pathogen interaction ^9, 10^. These genes are often upregulated in plants exposed to pathogen attack ^11, 12^, however in our *in vitro* seedling test, the transcript abundance of *ABCG23* rapidly decreased after inoculation with both *Alternaria* sp. and *Fusarium brachygibbosum* (Fig. 2e). Even though *F. brachygibbosum* is a major part of *N. attenuata’s* native pathosystem ^2^, the levels of *ABCG23* transcript did not differ among the tested lines after the inoculation with this pathogen, making it a less likely candidate to underlie the variation in survival rate observed in the field.

In contrast, the variation in *RLXL* transcript abundance observed among the tested lines after inoculation with both pathogens corresponded to differences in plant survival rates both *in vitro* and in the field (Fig. 2a, c, e). In our study, higher accumulation of *RLXL* transcripts was associated with lower seedling survival rates (as in UT-WT and P108), while the accessions with initially lower levels of *RLXL* transcript (as in AZ-WT, 106A, and P37) displayed better survival rates (Figs 2, 3). Therefore, *RLXL* appears beneficial when negatively regulated. The ribonucleolytic activity of *RLXL* does not seem to act directly on pathogen RNA, but rather indirectly by *e.g.,* degrading plants’ RNA and thereby modulating the expression of genes involved in plant response to the pathogen or triggering the production of small RNA molecules that contribute to the defense response ^13, 14^. Similarly, it could act as a signaling molecule in defense pathways, amplifying defense signals triggered by pathogen recognition receptors ^15, 16^. However, our analysis of transcript abundance of genes involved in signaling pathways of jasmonate, salicylic acid, and ethylene, commonly known to mediate plant response to pathogen attack ^4, 17, 18^ indicates that *RLXL* acts independently of hormonal signaling (Fig. S1).

While plants deploy RNases as part of their defense mechanism against infection, pathogens have evolved mechanisms to exploit these defenses for their own benefit ^19^. Common ways for pathogens to take advantage of plant RNases are *e.g.,* by utilizing small RNA fragments produced during host RNA degradation as signaling molecules to regulate their own gene expression or as a source of nutrients to facilitate their own growth ^20, 21^. Moreover, pathogens can directly manipulate the expression and activity of RNases to promote their own survival and proliferation ^22^ or produce RNA molecules that mimic host transcripts, allowing them to evade detection and degradation by host RNases ^23^. Regardless of initial *RLXL* transcription, all plants swiftly moved towards its suppression following the pathogen inoculation (Fig. 2e). Moreover, while pre-inoculation levels of *RLXL* in VIGS plants indicate successful knock-down (Fig. 4c), no detectable differences in *RLXL* transcript abundance were observed at the end of the experiment (Fig. 4f). Such findings align with the expected suppression of *RLXL* transcription in the presence of fungal pathogens. The retention of *RLXL* silencing in leaves at the conclusion of the experiment is reassuring (Fig. 4e). Considering that the leaves were not directly subjected to pathogen inoculation, there is no apparent reason for them to reduce their *RLXL* transcript abundance. During the course of our experiment, the co-inoculation with both pathogens seemed to predominantly impact the roots, leading to wilting symptoms that may arise from damage extending to the root-shoot junction ^24^. The suppression of *RLXL* occurs locally in the root tissues rather than on a whole-plant level. The lower initial levels of *RLXL* did not affect pathogenesis in the detached leaf assay, indicating that the *RLXL* effect might be specific to the root response to the pathosystem.

Our analysis of the link between *RLXL* transcript levels and seedling survival in an independent set of natural accessions showed that although AZ-WT and line P370 had higher survival rates, they did not exhibit significantly less disease symptoms (Figs 3, S2). Considering these results, the reduced mortality of the AZ-WT and P370 plants (and therefore the role of *RLXL* in the disease response) seems to stem from increased disease tolerance rather than resistance responses. We observed a decrease in the pathogenicity of our fungal lines between the first (Fig. 2) and the second (Fig. 3) *in vitro* trial. This variance could have been caused by the number of subculturing events of the pathogen cultures, or a different level of humidity in the Petri dishes of each trial. The difference in survival between UT-WT and AZ-WT was not replicated in the second trial after the inoculation with *Alternaria* sp. Instead, we observed a significant reduction in AZ-WT survival, which was not associated with a higher number of lesions (Fig. S2d), indicating that methodological factors may have caused this effect. Thus, in the second trial, we only draw conclusions from the *F. brachygibbosum* inoculation, but maintain that *RLXL* does not seem to be involved in the entry of pathogens into seedling tissues.

Although we have shown that reducing *RLXL* transcription contributes considerably to plant survival under pathogen attack, to fully understand the possible implications of *RLXL*-deficient plants within the ecosystem, it is necessary to look beyond individual plant to pathogen interactions. In ecological research, it has been well established that intraspecific diversity has a positive effect on the ecosystems, often leading to improved stability and productivity ^25–27^. Recent studies have revealed that the diversity of plant populations at a single locus (*i.e.*, allelic richness) can range from neutral ^28^ to negative effects ^29, 30^ on community productivity. This emphasizes the importance of investigating each effect individually, particularly in relation to different stress factors in communities.

Often, studies investigating the impact of allele richness on the performance of plant populations utilize binary designs with alleles represented in equal proportions ^27, 29^. However, a previous field study on *N. attenuata* plant populations showed that changes in the transcription of a single gene in 25% of field plants could cause dramatic changes in the yields of particular plants in population, altering the total yield gain ^31^. In the current study, we investigated four-plant subpopulations that differ in the frequency of the *RLXL*-deficient allele from 0% to 100% by increments of 25% (Fig. 6a). This design not only increased the variation in allele frequency, but also facilitated spatial analyses, enabling comparisons between plants with different alleles growing diagonally or adjacent to each other. We found that highest proportions of surviving plants in field populations occurred when the *RLXL*-deficiency allele (found in the AZ-WT accession – the ‘A’ allele) was present in one of the four plants in population (Fig. 6b). Additionally, the most beneficial position for the survival of *RLXL*-producing plants (with the *RLXL*-associated allele found in UT-WT accession – the ‘U’ allele) was diagonally across from the plant carrying the A allele (Fig. 6e). The survival probability of the U-plant in the diagonal position was around 50% higher than U-plants grown adjacent to an A-plant across the three field locations with low to medium disease intensity. The survival probability of a diagonal U-plant was not elevated if the A-plant died (Fig. 6e), indicating that the A-plants most likely do not act as dead-end pathogen sinks ^32^.

The populations with the beneficial A allele density were not only favorable for plant survival, but also for increased productivity. We observed a 14% increase in plant biomass (Fig. 6c) and an over 25% increase in seed capsule production (Fig. 6d) in the four-plant subpopulations with one A allele compared to populations with only the U allele. In agriculture, increasing diversity is a well-established strategy to enhance crop productivity, but in many cases its outcome is limited to 2-5% increase in yield ^33–35^. Therefore, even though the differences in productivity observed here represent small and medium effect sizes due to replicate availability (Table S2), they are still several orders of magnitude higher than what is usually expected. Our results support the findings that increasing biodiversity alone might not be enough to improve survival and productivity during disease outbreaks in agro- or natural ecosystems. Although the movement of pathogens around populations leading to overall benefits has been theorized for many years ^36, 37^, there is still little experimental evidence for mechanisms of frequency- and spatially-dependent effects, especially when conferred from single loci. Our study offers experimental evidence to support this perspective. However, to strengthen our conclusions, further research involving genetic modification of *RLXL* transcription under field conditions, with increased replicates of modified plants at different frequencies and in varying positions in population, is warranted. Mechanisms of plant community resistance to a fungal pathogen described here presents a great tool for improving current conservation or agricultural models and facilitating the generation of modern cultivar mixtures that promote positive interactions between the plants ^38^.

## Methods

### Biological material and growth conditions

Four replicates of a previously described *Nicotiana attenuata* advanced intercross - recombinant inbred line (AI-RIL) population ^7, 39, 40^ were planted in the field in 2017 (Lytle Ranch Preserve, Santa Clara, Utah; ‘Snow plot’, N37.141283, W114.027620). Plant germination and field adaptation were carried out as described previously ^41^. AI-RIL (F12 generation) replicates were randomly distributed across the field site, within four-plant subpopulations planted around individual emitters of a drip irrigation system. Each subpopulation included one Utah wildtype control (UT-WT, 31x inbred ^39^) planted in one of the four spots, and three randomly selected AI-RILs. The watering system was turned on for 1 hour in the morning and evening until established, and then as needed. The founder lines of the AI-RIL population, UT-WT and Arizona wildtype (AZ-WT, 21x inbred ^40^) as well as AI-RIL 106A, which showed high resistance to the *Fusarium-Alternaria* pathosystem, were grown in the Snow plot the following year (2018) and monitored for health and survival in the face of the recurrent pathosystem. Plant growth and monitoring was as described previously for the focus plants (n = 15-58; ^6^). Natural accessions P108, P370, and P422 used in the *in vitro* seedling experiment (Fig. 3, Fig. S2) were selected based on their genetic information (F2 generation, ^42, 43^). Seeds for all *in vitro* tests (Fig. 2, Fig. 3, Fig. S1, Fig. S2) were germinated as previously described ^44^ and plated in a circle of eight in each dish with the center area left empty for a fungal plug. Additionally, *in vitro* cultures of two fungal stains, *Fusarium brachygibbosum* Padwick Utah 4 and *Alternaria* sp. Utah 10 were used in this study. The *in vitro* maintenance and the culture conditions were described previously ^2, 4, 6^. Both strains were re-isolated from diseased seedlings before the first *in vitro* seedling test.

### 2017 Field experiment

The mortality of the AI-RIL population field experiment was assessed 70 days post planting, right before the harvest (Fig. 1a). Dying plants had a characteristic black or brown discoloration of more than 1 cm at the bottom of the stem, as well as wilting leaves and apical meristems. Plants which exhibited these symptoms or were removed previously due to vasculature failure and thus stem collapse were characterized as dead. The resistance scores for each AI-RIL were calculated based on four factors: 1. whether the AI-RIL itself was dead or alive, 2. whether the UT-WT control within the four-plant subpopulation was dead or alive, 3. total number of plants that remained alive within the subpopulation, and 4. the location of the subpopulation in the field plot (Fig. 1c). The second factor modifies the impact of the first factor. For example, an AI-RIL that survived while its UT-WT control died should have a higher resistance score than if the UT-WT had survived. The third factor further tunes the resistance of the AI-RIL to the number of other plants that survived in its group: if the AI-RIL is the only one to survive, it should have a higher resistance score, because it survived not only the pathosystem above all its neighbors, but also a high pathosystem prevalence. Finally, there was a clear increase in pathosystem presence from one side of the field to the other (Fig. 1a), and thus an environmental correction was applied in the resistance scores. The whole of the irrigation system was divided into four sections, from left to right (lines 1 – 4, Fig. 1a, Fig. 6), of equal width across the field. The last section to the right was not completely filled with plants. Sections 2 through 4 demonstrated a linear increase in pathosystem presence and mortality. If an AI-RIL survived in section 4, for instance, its resistance score was inflated by 4. Similarly, resistance score of surviving plants in lines 1-3 were inflated by 1-3, respectively. The resistance scores of the four replicates of each AI-RIL were added together to produce the overall resistance score for that AI-RIL line. UT-WT controls and any AI-RIL that did not have four replicates due to early, non-pathosystem-related deaths were excluded, as the resistance sum could be biased by the number of replicates. The final, even distribution of resistance scores across the field is shown (Fig. 1d).

### Quantitative trait locus (QTL) mapping

The genotype information of the AI-RIL population and the linkage map were described previously ^7, 45^. The final resistance score of all the AI-RILs was tested for associations with particular SNPs in the *N. attenuata* genome (manually coded QTL mapping, 100 bootstraps, Fig. 1e). Prior to bootstrapping, the resistance scores were log normalized to meet the assumption of normal residuals required for parametric QTL mapping. The strength of potential association of each SNP to the phenotype is reported with a logarithm of odds (LOD) score. A threshold LOD for determining the significance of a particular SNP association was calculated by running the model with random RIL scores assigned to each RIL SNP profile. Any SNP above the LOD threshold has a less than 5% chance of its potential association being due to randomness (95% confidence interval).

### In vitro seedling tests

AZ-WT, UT-WT, and AI-RIL line 106A (n = 7-9 plates of eight seedlings, Fig. 2, Fig. S1) as well as a five different natural accessions of *N. attenuata* (n = 6-9 plates of eight seedlings, Fig. 3, Fig. S2) were grown in Petri dishes for follow up analyses. Two weeks post germination a subset of plates from each genotype was harvested destructively and flash frozen to serve as a control (the “0” hour samples). Seedlings from each plate were pooled as one replicate. The remaining plates were inoculated with fungal plugs (5 mm in diameter) containing either *F. brachygibbosum* or *Alternaria* sp. culture. Fungal plugs were taken from concentric ring at the same distance from the center of fungal source plate to ensure similar age of the fungus and applied to the center of the seedling plate depending on its treatment. Fungal plugs from each source plate were distributed across treatment groups. Plates were rewrapped and placed back into the incubator until 1-, 6-, or 12-hours post inoculation at which time subsets of the remaining replicates were sampled in the same manner as the controls. After the 12-hour post inoculation time point, 6-9 plates of each treatment remained in the incubator and were left undisturbed until 15 days post inoculation (dpi), at which time they were evaluated for their mortality. Seedlings that were marked as dead were translucent and had collapsed. Those that had survived (were not translucent or collapsed) were scored for visible lesions.

### Transcript Abundance – qRT-PCR

Either entire seedlings (Fig. 2D,E, Fig. 3B, Fig. S1) or leaf (150 mg) and root tissue (300 mg; Fig. 4) were flash frozen in 2 ml Eppendorf tubes, and extracted for RNA with TRIzol reagent (Invitrogen) according to the manufacturer’s instructions. Total RNA was quantified using a NanoDrop (Thermo Scientific, Wilmington, DE, USA) and cDNA was synthesized from 500 ng of total RNA using RevertAid H Minus reverse transcriptase (Fermentas, Vilnius, Lithuania) and oligo (dT) primer (Fermentas). Quantitative real time PCR (qRT-PCR) was performed in an Mx3005P PCR cycler (Stratagene, San Diego, CA, USA) using a 5X Takyon for Probe Assay (no ROX) Kit (Eurogentec, Liège, Belgium) with TaqMan primer pairs and a double fluorescent dye-labeled probe for testing the silencing efficiency of the intracellular ribonuclease LX-like (*RLXL*) (Fig 4C). *N. attenuata’s* sulfite reductase (*NaECI*) was used as a housekeeping gene, as described previously ^46^. For all other transcript abundance analyses, a SYBR green reaction mix (with ROX; Eurogentec, Liège, Belgium) was used. The sequences of primers and probes used for qRT-PCR are provided in Table S1. All qRT-PCR data were normalized using the delta-Ct method.

### Virus Induced Gene Silencing (VIGS)

A single gene from the region of interest, intracellular ribonuclease LX-like (*RLXL,* NIATv7_g12084, Table S2), was transiently silenced using VIGS in a background of UT-WT and P-108 as described previously ^47, 48^. Briefly, a PCR with primer pair *RLXL*-1F (5’-GCGGCGGTCGACCAGAAGATTTCTTATTTCCAAG-3’) and *RLXL*-1R (5’-GCGGCGGGATCCCTTTACATCTCTTACTTTCTGG-3’) and cDNA from *N. attenuata* roots harvested 2 h after leaf treatment with *Manduca sexta* regurgitant as template was performed. The resulting 301 bp PCR fragment was then digested with *Sal*I and *Bam*HI and cloned in vector pTV00 cut with the same enzymes, resulting in the *NaRLXL* silencing vector *pTV::RLXL* (5.8 kb). These were compared to VIGS plants with an empty-vector (*EV*) construct in their resistance to both a leaf and root inoculation with *F. brachygibbosum*. Additionally, the *pTV::PDS5* vector, harboring a part of a phytoene desaturase (*PDS*) which causes extensive leaf bleaching (Fig. 4a), was used to monitor the progress of gene silencing. Plants for this experiment were germinated and grown as described previously ^44^ up to the TEKU stage (∼20 dpg) when they were transferred to pots in isolated climate chambers (26°C, 65% relative humidity, 16 h day : 8 h night light cycle; n = 10 per treatment). They were infiltrated with the silencing construct specific to their treatment within one week after the transfer and continued to grow until the first occurrence of leaf bleaching on the *PDS* control plants. A subset of the VIGS plants was sampled destructively for root tissue 21 days after VIGS (DAV) to determine the rate of successful silencing (Fig. 4).

### Detached leaf inoculation assay

To avoid any influence from and variation caused by mechanical leaf damage a leaf inoculation assay was performed on detached leaves as described previously ^2, 4, 6^. Briefly, the “+1”, “+2” and “+3” leaves ^49^ of VIGS plants were removed 23 DAV and placed, abaxial side up, in square Petri dishes with four layers of moist autoclaved tissue paper. Fungal mycelial plugs (3 mm diameter) were cut from the marginal regions of 14 day-old actively-growing *F. brachygibbosum* cultures and placed on the abaxial sides of the leaves. Disease symptoms were assessed six days later by determining the lesion size in 3-4 independent biological replicates (3 leaves per biological replicate) per genotype, six lesions per leaf.

### Root Dip Assay

The VIGS plants were inoculated with *F. brachygibbosum* cultures using a protocol previously utilized on 10 and 20 day old seedlings ^2^ adapted to facilitate the inoculation of 49 day old plants. Briefly, 24 DAV the roots of the VIGS plants were dipped by placing pots in a pot tray with concentrated (>10^5^ spores mL^-1^) solutions of *F. brachygibbosum* for 30 seconds before returning them to their original individual pot trays (Fig. 4A). Relative growth rate (RGR) for elongation was measured before and after root-dip inoculation at 0, 15-, and 23-days post inoculation (dpi). The percentage of leaf wilting was estimated at 15 and 23 dpi by counting the number of visibly wilted leaves out of the total number of leaves per individual at each time point. Additionally, images of each plant were taken for subsequent analysis of the leaf angle (Fig. 5). Finally, leaf and root samples were collected destructively 23 dpi (47 DAV) at the conclusion of the experiment and flash frozen for further analyses (Fig. 4e,f).

### Image processing

All images were taken from the same distance and angle from the VIGS pots and processed in Fiji (ImageJ2, Version 2.3.0/1.53f) using the length and angle measurement tools. Leaf length was taken from the attachment point of the leaf with the stem to the tip of the leaf, despite leaf curvature. Leaf angle was taken between this leaf length measurement line and the horizontal axis. Measurements were taken on 3 randomly chosen leaves per picture.

### Statistical analysis

All data were analyzed using R version 4.1.1 ^50^ and RStudio version 1.3.1073 ^51^. Most datasets were fit to a linear model and checked for homoscedasticity and normality (through a graphical analysis of residuals) ^52^. Outliers were removed only after identification through an evaluation of Cook’s distance and leverage. Pairwise *post hoc* comparisons were extracted per the contrasts tested (between all pairs of bars in respective panels, or between all pairs of points per timepoint, in Fig. 2b,d,e, Fig. 3, Fig. 4c,e,f, Fig. 5b-e, Fig. S1, and Fig. S2; *emmeans* ^53^), after significance of fixed effects in ANOVAs.

The mortality analysis on the 2018 field experiment data (Fig. 2a) was performed using a multivariate Cox regression (*coxph()* function, *survival* package in R ^54, 55^). The distribution of survival times across genotypes was visualized using the function *survfit()*.

Finally, due to the variable nature of field data originating from complex population dynamics, ANOVAs found no significances for Fig. 6b-d, though clear trends could be observed. Glass’s effect size is often used in field measurements to evaluate the importance of changes that are visually, but not statistically observed; this Δ calculates proportions of each treatment group above and below the mean of the control group, normalized to the control’s standard deviation ^56^. For Glass’s Δ values, a small effect size is considered to be 0.20-0.35, and more notably, a medium effect size is considered from 0.35-0.65 and a large one above 0.65 (Fig. 6c,d). For Fig. 6e, an ANOVA produced a significant result by the position of the plant relative to the present or absent *RLXL*-deficient (A) or -producing (U) plant in the population.

## Acknowledgements

We thank the glasshouse department at the Max Planck Institute for Chemical Ecology and the field teams in 2017 and 2018 for their support; Brigham Young University for the use of the Lytle Ranch Preserve field station in Utah, USA, and APHIS for constructive regulatory oversight; the technical staff at the department of Molecular Ecology for providing seeds; both the International Max Planck Research School (IMPRS) on the Exploration of Ecological Interactions with Chemical and Molecular Techniques and the Young Biodiversity Research Training Group (yDiv) for their support of HFV and EM.

## Author contributions

Conceptualization: EM, HFV; experimental investigation: EM, HFV, PB, MP, KG; data analysis: EM, HFV, PB; writing – original draft: PB; writing – review and editing: HFV, EM, KG, MP, ITB; resources: ITB; visualization: EM, HFV, PB.

## Notes

### Competing Interest Statement

The authors have declared no competing interest.

