## Supplemental files for "QTL mapping in field plant populations reveals a genetic basis for frequency- and spatially-specific fungal pathosystem resistance"

**Figure S1: Common plant hormone pathways do not explain differences in plant resistance to *Alternaria* sp. and *Fusarium brachygibbosum* in an *RLXL*-dependent manner.**

The analysis of relative transcript abundance of genes related to (a) jasmonate, (b) salicylic acid, and (c) ethylene signaling at 0-, 1-, 6- and 12-hours post inoculation with either *Alternaria* sp. or *Fusarium brachygibbosum*. Lowercase letters indicate statistically significant differences between the genotypes within one timepoint based on ANOVA followed by Tukey adjusted pairwise contrasts (± SE, n = 6 - 9 per timepoint).

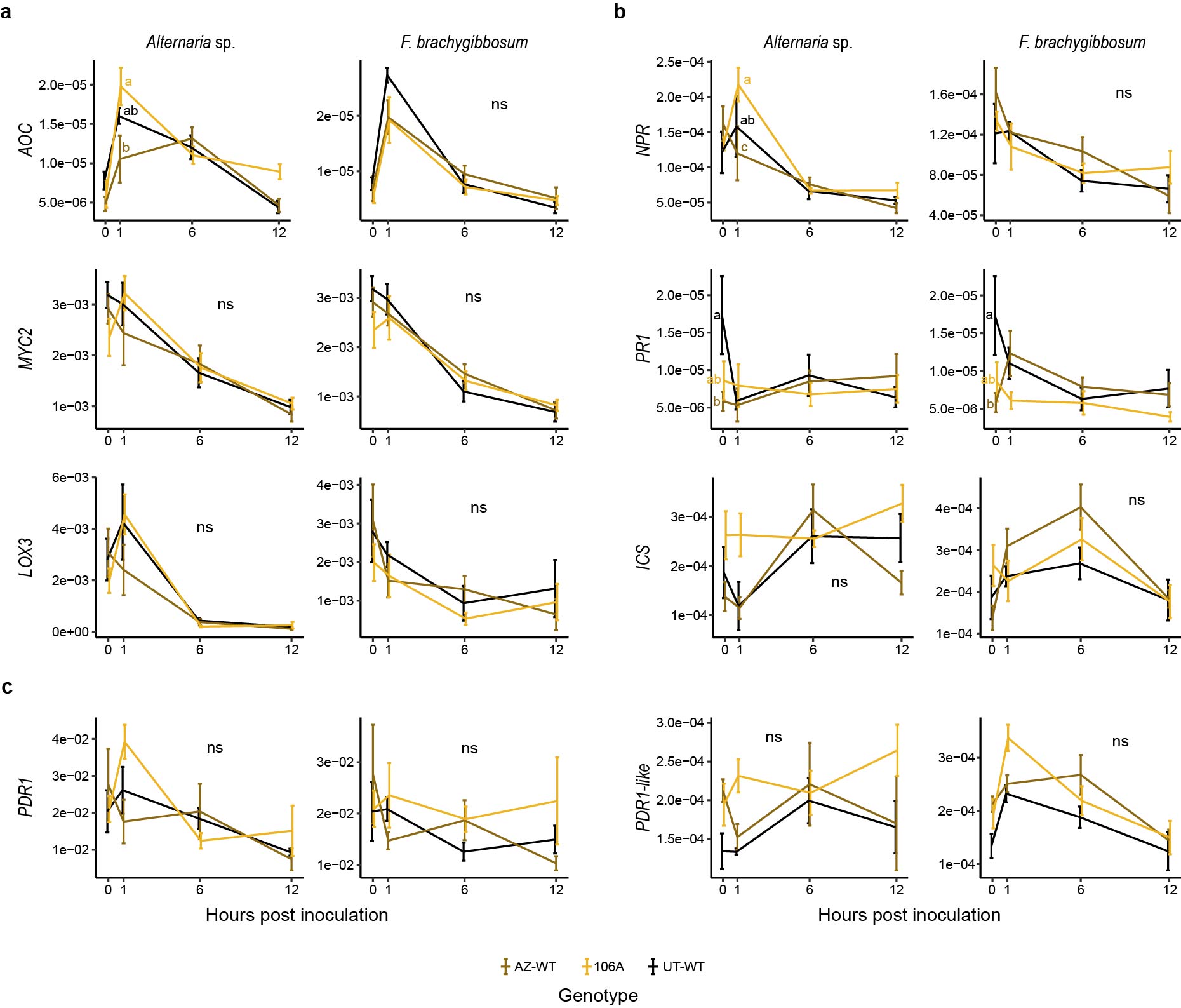

**Figure S2: Variation in seedling survival phenotypes among natural accessions of *Nicotiana attenuata* after the inoculation with *Fusarium brachygibbosum* or *Alternaria* sp.**

(a) Proportion of seedlings without damage, and (b) seedlings with lesions 15 days after inoculation with *F. brachygibbosum*, or proportion (c) of seedlings without damage, (d) seedlings with lesions, and (e) seedlings surviving 15 days after inoculation with *Alternaria* sp. Lowercase letters indicate statistically significant differences based on ANOVA followed by Tukey adjusted pairwise contrasts (± SE, n = 6-9 per accession).

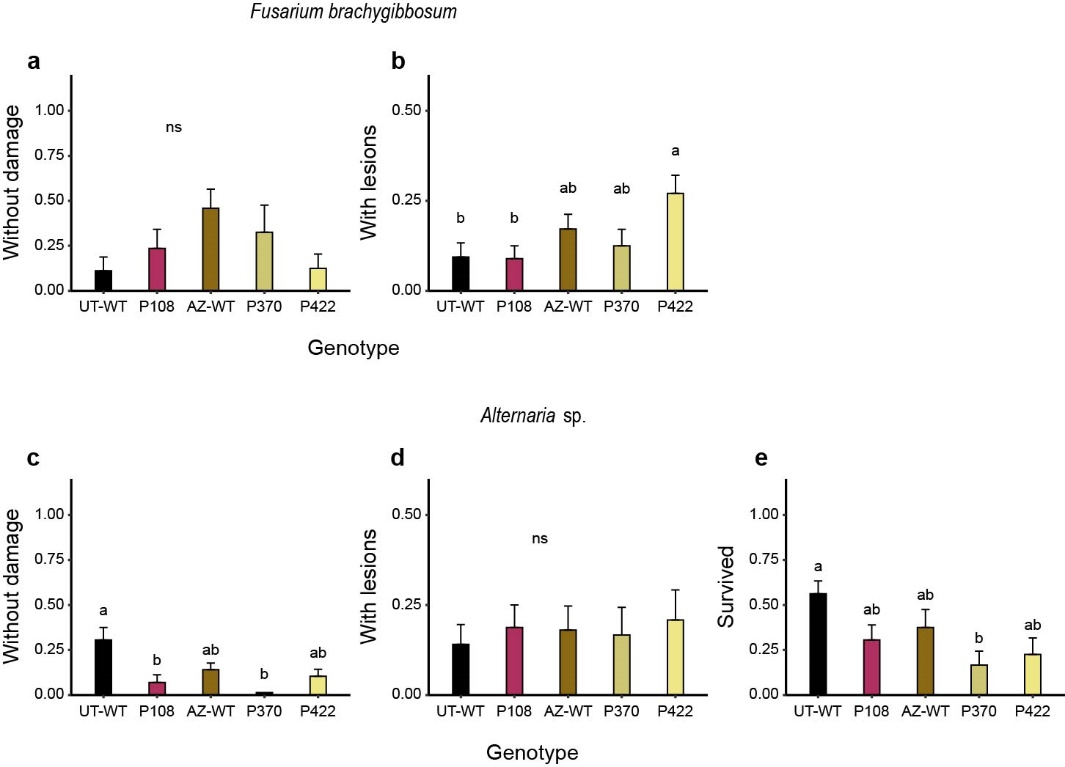

**Table S1: Candidate genes potentially involved in plant pathogen resistance.**

The functional annotation and gene expression profiles are based on *N. attenuata* NADH browser (Brockmoeller et al. 2017). The genes selected for further characterization are shown in bold.

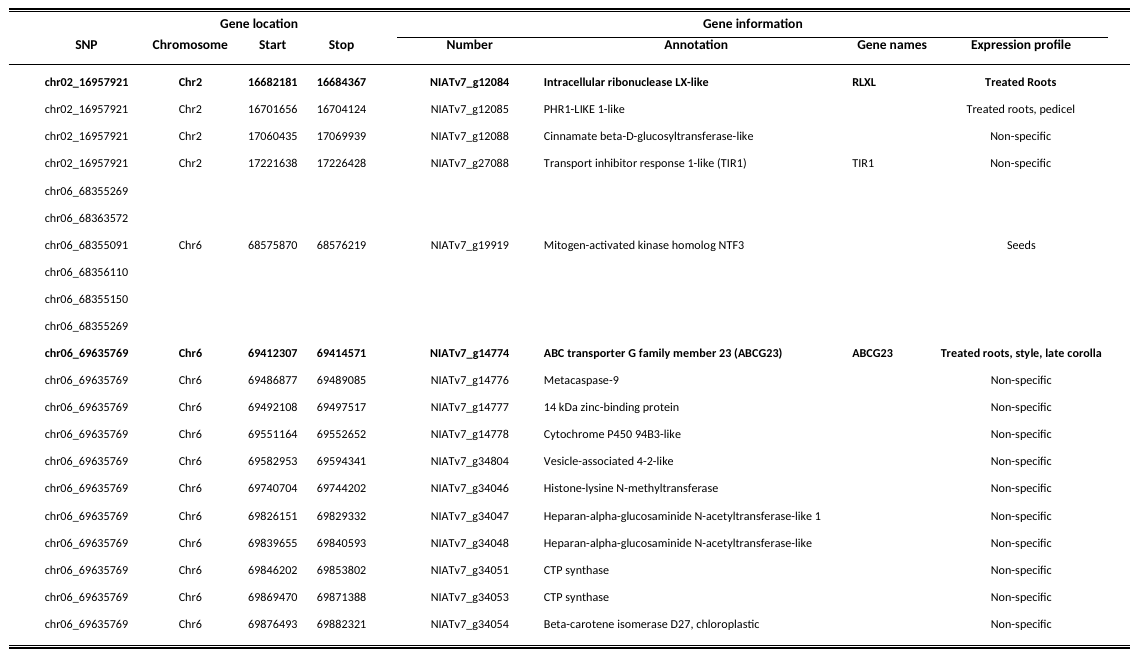

**Table S2: Statistical summary of the biomass accumulation and seed capsules number for populations with one AZ-WT-like RLXL allele (dead or alive) or populations with no AZ-WT-like RLXL alleles with all plants alive or one plant dead.**

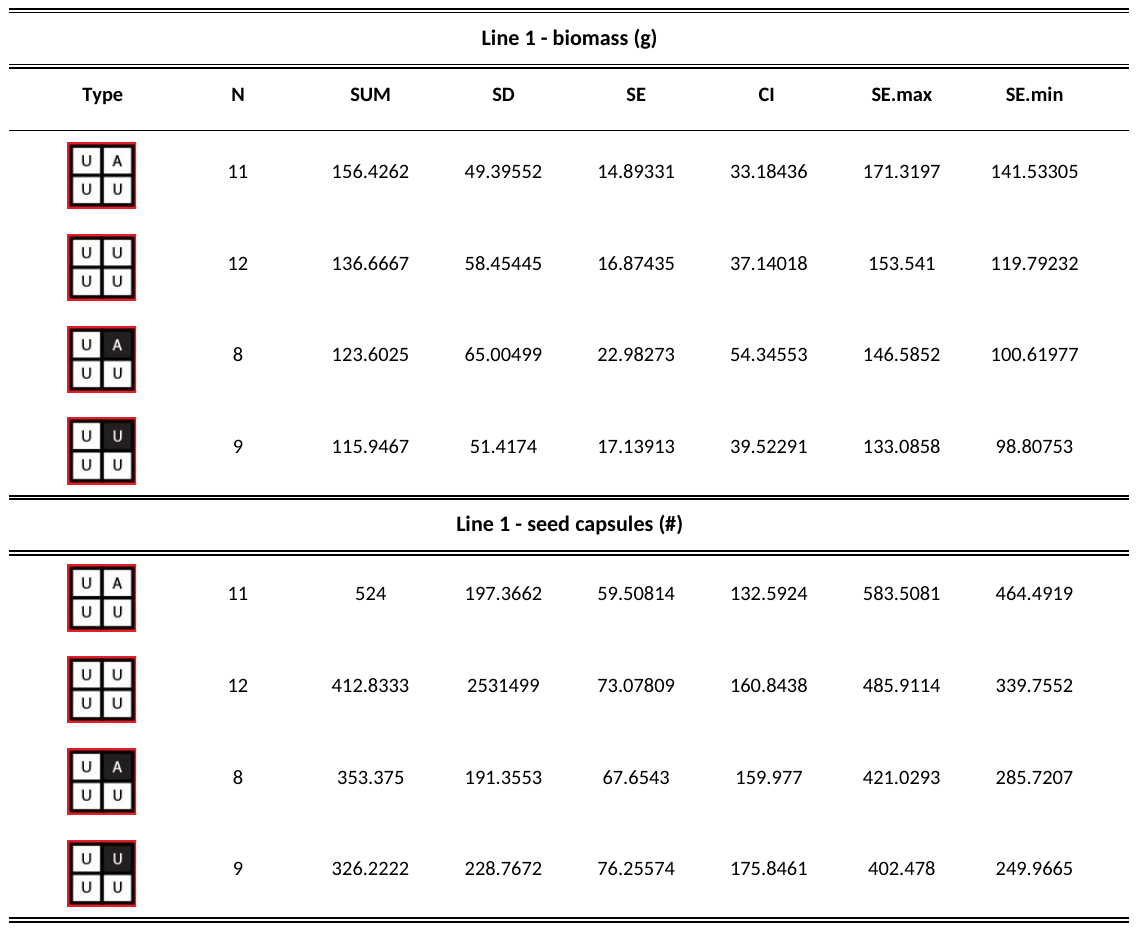

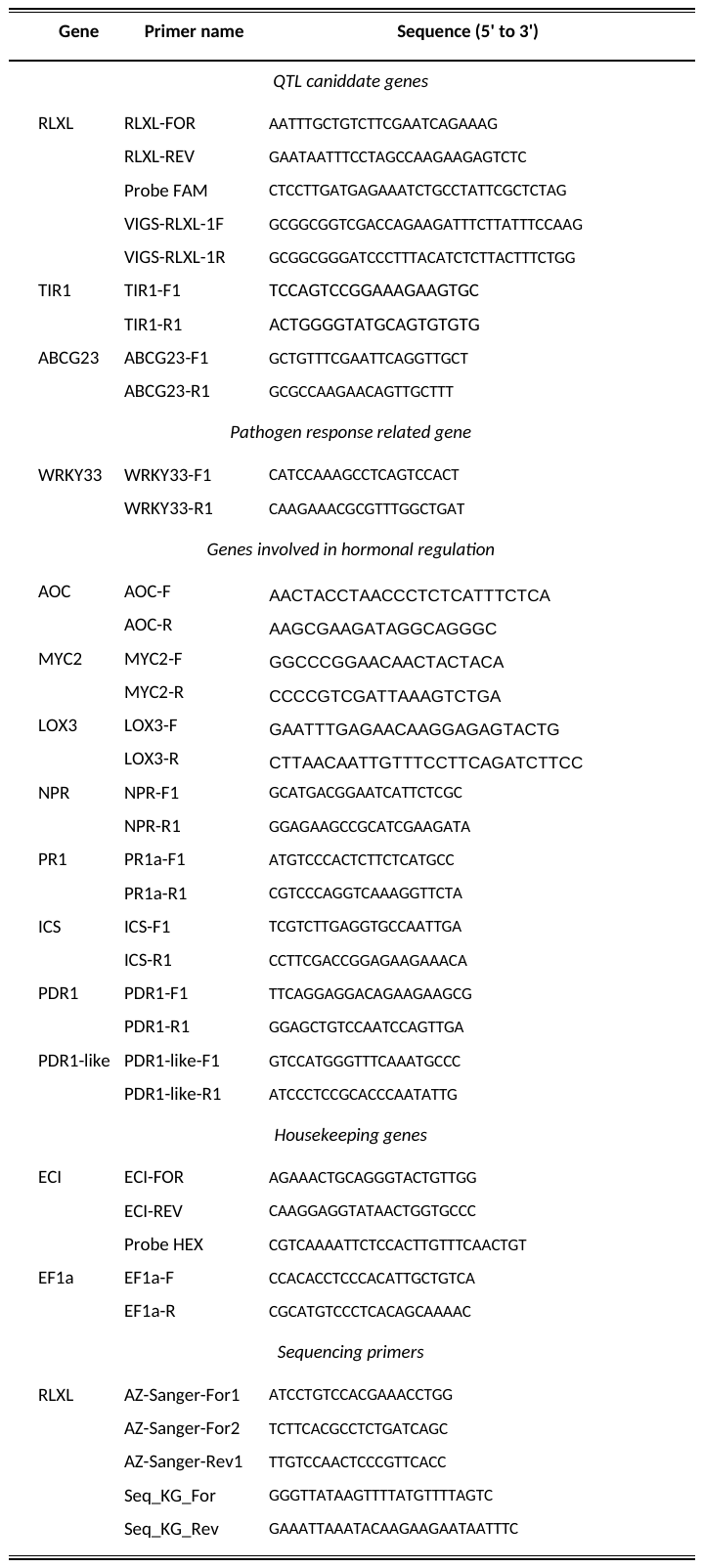
**Table S3: Primer sequences used in this study**
